# Modelling Human Gait using a Nonlinear Differential Equation

**DOI:** 10.1101/2021.03.16.435713

**Authors:** Jelena Schmalz, David Paul, Kathleen Shorter, Xenia Schmalz, Matthew Cooper, Aron Murphy

## Abstract

We introduce an innovative method for the investigation of human gait, which is based on the visualisation of the vertical component of the movement of the centre of mass during walking or running, in the space of the coordinates position, velocity, and acceleration of the centre of mass. Collected data has been numerically approximated by the best fitting curve for a non-linear model. The resulting equation for the best fitting plane or curve in this space is a differential equation of second order. The model that we suggest is a Duffing equation with coefficients that depend on the height of a walker or runner and on the angular frequency of the oscillation. Statistics about the distribution of the Duffing stiffness depending on the speed is presented.

**Author Summary:** We study the human gait modelled by the movement of the centre of mass of the test person. This is an example of a biological process which can be considered as a periodical dynamic system. Roughly, this movement behaves in a similar way to a vibrating mass suspended on a spring, but it is more complex. The vertical component of the movement during walking or running can be visualised as an oscillogram: a graph of the position as a function of time. We suggest a visualisation of the data in 3D space, where the coordinates describe position, velocity, and acceleration. Our new visualisation method allows us to model the movement of a person’s centre of mass by a nonlinear differential equation. The resulting curve for an ideal spring-mass movement, without viscosity or external force, is an ellipse in the suggested 3D space. The shape of the data curve shows at which position an additional force was applied, or the movement slowed down. Some deviations are common for all test persons and others are different. In the future we plan to investigate the reasons for these deviations, such as different running techniques or the presence of injuries.

## 2 Introduction

### 2.1 Background

Research on the mechanics of human gait can be of interest to different disciplines, for example sport science, medicine, and robotics. In this paper we discuss a model for the movement of the vertical coordinate of a person’s centre of mass (COM) during walking and running.

Human locomotion is an inherently complicated process requiring the complex integration of neural and musculoskeletal control in response to both internal and external forces. In an attempt to strip away complexity and gain an understanding of the fundamental principles underpinning human locomotion, simple mechanical models have been developed. The mechanical simplification of locomotion allows the identification of just a few key parameters that can be manipulated to examine cause and effect relationships and identify which features most influence the system [4, 20].

Blickhan suggested a linear spring-mass model for hopping in 1989, [1]. Other papers followed, for example [2, 9, 24, 25]. The motion of the centre of mass is described by the equation *mz*_*tt*_ + *Kz* = *−mg*, where *m* is the body mass, *z* is the vertical deflection of the centre of mass with the origin on the treadmill surface and the direction chosen upwards. The constant *K* is the stiffness, and *g* is gravitational acceleration. By *z*_*tt*_ we denote the second derivative of *z*, i.e. the vertical acceleration of the centre of mass.

There have been different approaches on how to calculate leg stiffness. Blickhan’s approach uses the formula 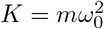, where *ω*_0_ is the stride’s angular frequency of the oscillation, which, during gait, reflects the stride’s angular frequency. [1, 24].

The other approach for calculation of the leg stiffness is to find the ratio of *F*_*max*_, the maximum value of the vertical ground reaction force, and Δ*L*, the absolute value of the leg compression, i.e. 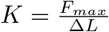. This definition of leg stiffness is used in several papers [2,5, 8,9, 25]. An overview of these two approaches is presented in [24]. There is a third approach to leg stiffness calculation based on the measurements of loss of mechanical energy by walking/running, *W* . Leg stiffness *K* is found from the formula 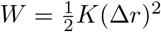, where Δ*r* is the shortening of a spring (e.g. [6]).

In examining mechanical and metabolic determinants of the human walking gait, Kuo [19], and [7] employed an anthropomorphic three-dimensional, passive-dynamic model, in which human legs were represented as rigid inverted pendulums with small point masses modelling each foot and a larger mass modelling the concentration of the COM at the pelvis. These studies drew on earlier models of a rigid swing leg during walking [23] and continued the view that walking and running were two distinct gaits that could not be described using similar mechanical models. This view, however, was discounted by Geyer, Seyfarth, Blickhan (2006) [12] who demonstrated that a compliant-legged, spring-mass bipedal model consisting of two linear, equal and massless springs and a single COM point mass, as an extension of Blickhan’s one-dimensional model, reliably predicted ground reaction forces and COM behaviour in both human walking and running. Several subsequent studies further validated the efficacy of a bipedal spring-mass model of walking [14,17,18,20,26]. While much of the twenty-first century research in the field has adopted the bipedal spring-mass model and focused on adapting or adding selected elements to improve prediction accuracy for both walking and running gait mechanics, Blickhan’s spring-mass model remains largely valid and has been applied, with modifications to suit certain parameters, in recent studies [21, 22].

Our mathematical model is based on the analysis of the collected data of the three dimensional movement of COM. In this paper we concentrate on the projection of the movement of COM on the vertical axis. We plan to collect more data and apply our new method for the investigation of projections on the horizontal axes – the axis along the walking/running movement and the axis that is perpendicular to it, see Section 9.

We suggest a new approach for finding leg stiffness for the simple harmonic oscillation model, and then develop a more precise model assuming that the stiffness for a fixed speed is not a constant but depends on the displacement of the COM. The innovative idea of our method is to visualise the data for the vertical component of a motion of the COM, *z*(*t*), as a curve in the three dimensional space, (*z, z*_*t*_, *z*_*tt*_) (see Section 3). Here *z*_*t*_ and *z*_*tt*_ are the first and the second derivatives of the function *z*(*t*), correspondingly. These curves are interesting by themselves as they are individual for each person and velocity. We provide examples of the variety of these curves in Section 9. In this paper we discuss the properties that are common for these curves. We apply the Fast Fourier Transform (FFT) to the data and filter out the smaller frequencies.

## 3 Data Recording

We have collected data for six participants aged from 18 to 55 years, three males and three females. The study followed ethical protocols as per ethics requirements (HE19-239). We have measured the vertical coordinates *z*(*t*) of the COM for each participant walking or running on the treadmill. The markers we used were the Left and Right PSIS and ASIS, then we computed the average of all four. The data was collected for different integer velocities, at 100 frames per second, over 10 seconds, for each velocity, using an 8 camera, Qualisys Motion capture system with the COM reconstructed using a pelvic marker set within Visual3D.

Using MATLAB, we have visualised the data as curves in the three-dimensional space (*z, z*_*t*_, *z*_*tt*_). We have filtered out high frequency oscillations, such as noise and individual features, using the fast Fourier transform function in MATLAB. For FFT threshold equal to 0.3, the curves are close to an ellipse. The direction of the motion along the curve is found, for example, in the following way. Find the point with the greatest *z*-coordinate. This is the highest point of the centre of mass during the gait cycle. The velocity at this point is equal to zero. Hereafter the movement goes down, i.e. the velocity *z*_*t*_ becomes negative. In both pictures of Figure 1, from those perspectives, the movement is clockwise.

**Fig 1.**
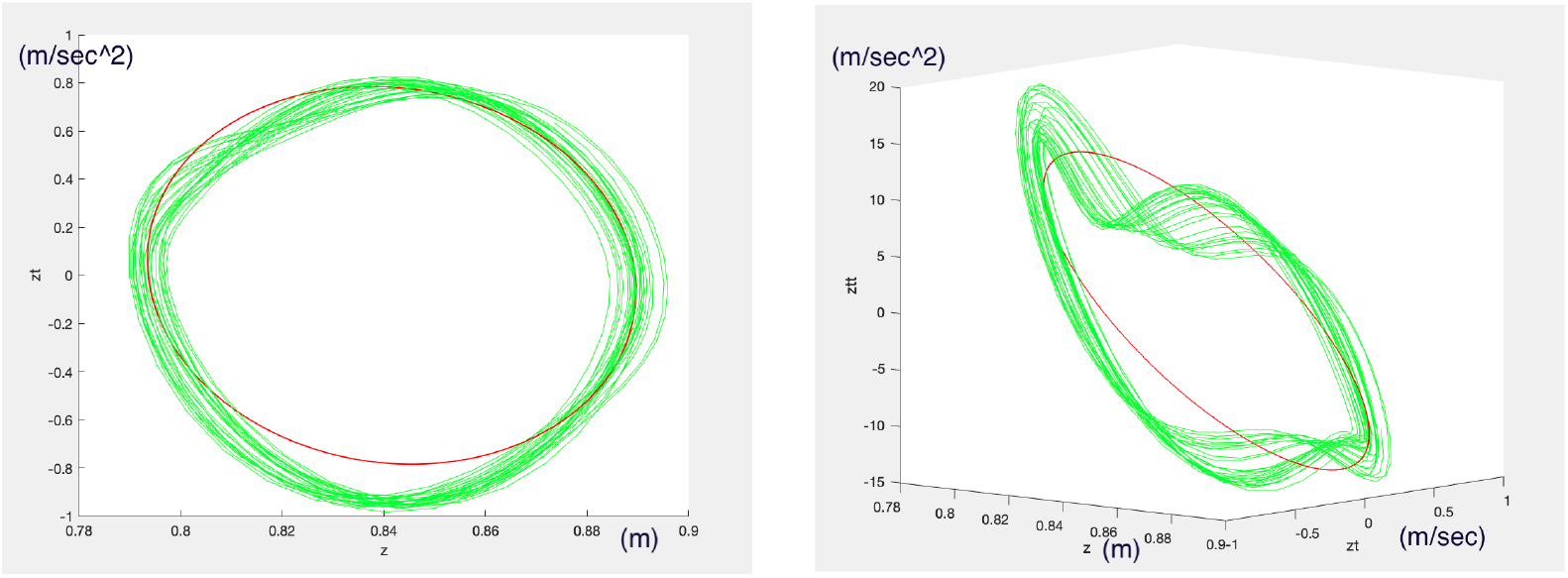
A MATLAB 3D figure shown from different perspectives. The green curve is a smoothed data curve with high-pass FFT threshold 0.03, the red curve is a smoothed data with FFT threshold 0.3.

## 4 Data Interpretation

The pictures in the space with the coordinates position, velocity, and acceleration, are rich in information. For example, Figures 2 and 3 show the data for walking (4km/hr) and running (9km/hr) of a participant, correspondingly. Here we look into the projection to the plane *z, z*_*tt*_, i.e. the horizontal axis shows the position of the COM (in meters) and the vertical axis shows the acceleration (in *m/sec*^2^). The part CD of both curves corresponds to the phase of the gait when a foot touches the surface of the treadmill. In the case of walking it is a flat line; vertical acceleration is close to zero. In the case of running, acceleration is diminishing because of braking during initial foot contact. The phase DE corresponds to the propulsion during toe-off, where acceleration (and consequently force) increases. On the segment EA acceleration diminishes, turns to zero when the the COM reaches its average position, and is minimal at the point A. The minimum acceleration for the walking curve is *−*2 *m/sec*^2^, the minimum acceleration for the running curve is about *−*10 *m/sec*^2^, i.e. close to the gravitation constant *g*. The acceleration is less during walking than during running because the body is always in contact with the ground whereas during running there is a flight phase. The arc AB on Figure 3 corresponds to the flight phase of running. This part of the curve is more complicated than just a flat constant *z*_*tt*_ = *−g*, because it is smoothly connected with the rest of the curve. Some information that we get from these curves is common for all participants and walking/running speeds, but some features are individual – for example not each participant has the “flight” component AB at running speeds, due to individualised transitions between walking and running gait patterns. Our aim in the future is to collect data for more participants and compare individual properties of the curves, see Section 9. In this paper we concentrate on their common properties, and suggest three models, based on differential equations. We build our three models based on data, purely numerically and mathematically. The first model is well-known, it is a harmonic oscillation. Our novelty here is the method we develop for computing leg stiffness.

**Fig 2.**
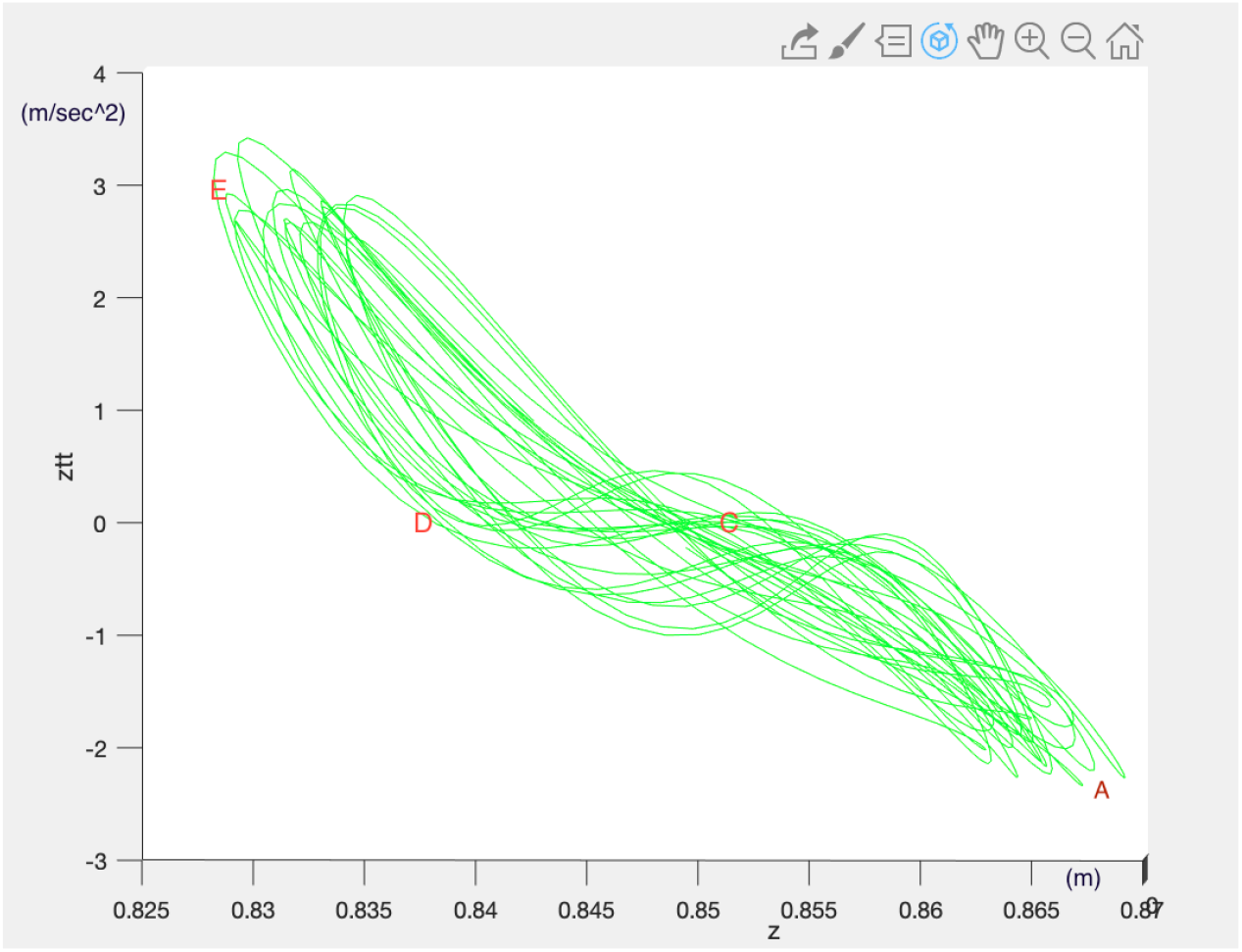
Data for walking, 4km/hr. The horizontal axis shows the position and the vertical axis shows the acceleration of the COM. The part CD corresponds to the phase of the gait when a foot touches the surface, DE corresponds to the propulsion during toe-off, during EA the COM moves upwards and the acceleration diminishes.

**Fig 3.**
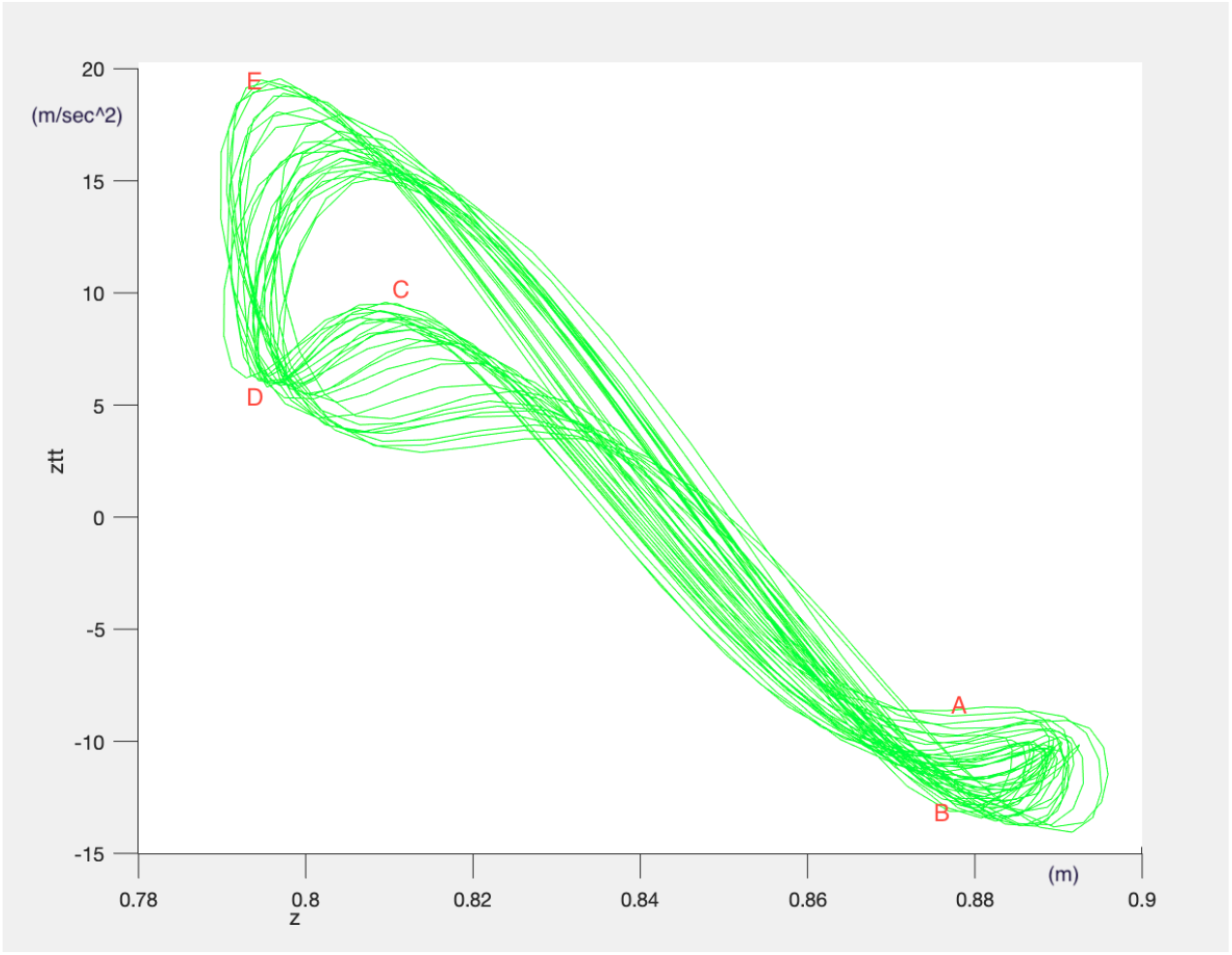
Data for running, 9km/hr. The horizontal axis shows the position and the vertical axis shows the acceleration of the COM. The part CD corresponds to the phase of the gait when a foot touches the surface, DE corresponds to the propulsion during toe-off, during EA the COM moves upwards and the acceleration diminishes. The arc AB corresponds to the flight phase.

## 5 Modelling Gait as Harmonic Oscillator

### 5.1 Best fitting plane and the interpretation of its coefficients

We begin with visualisation of the data. Consider a curve in the three dimensional space (*z, z*_*t*_, *z*_*tt*_) that represents smoothed gait data, as for instance the red line on Figure 1 (FFT=0.3). This curve is not quite flat, but it is close to a flat ellipse with the centre at *z* = *z*_0_, *z*_*t*_ = *z*_*tt*_ = 0. The equation of a plane in three dimensional space (*z, z*_*t*_, *z*_*tt*_) that passes through the point (*z*_0_, 0, 0) is

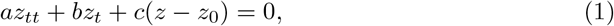

where *a, b*, and *c* are constants, and *z − z*_0_ is the vertical displacement of the average of the centre of mass. The zero for *z*(*t*) is chosen to be on the level of the treadmill, and the *z*-axis direction is upwards.

We find the coefficients *a, b*, and *c* in Eq. (1) numerically, by finding the best fitting plane for the smoothed data curve. The equation for the plane is interpreted as a second order differential equation that describes a damped driven harmonic oscillator. The coefficients in Eq. (1), in the context of the spring model, have the meaning of *a* = *m*, which is the mass of the oscillating object, *b* = *ν*, which is the viscosity of the spring, and *c* = *K*, which is the spring stiffness,

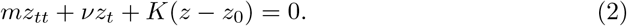

If we consider a stable gait, we need to assume, in this linear model, that the viscosity is zero. Eq. (2) turns to

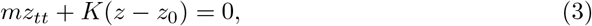

which is Hook’s Law [15] that describes a movement of a spring with stiffness *K*.

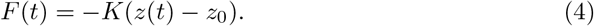

### 5.2 Where the gravitation constant is hidden

We note that the gravitation constant is included in Eq. (2) implicitly. Namely, we can rewrite the equation as

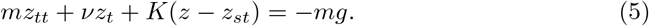

The coordinate *z*_*st*_ is the average of the vertical coordinate of the centre of mass of a standing body but in a walking/running posture. It is not exactly the same as the coordinate of COM in a standing position. The relation between *z*_0_ and *z*_*st*_ is calculated from Eqs. (2) and (5) and is 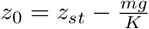.

## 6 Modelling by a non-linear homogeneous differential equation

### 6.1 Best fitting curve, interpreted as a non-linear second order differential equation

Non-linear gait dynamics has been discussed, for example, in [13]. The author remarks that stride-to-stride fluctuations, which are often considered to be noise, actually convey important information. With the aim to describe these fluctuations, we refine the method used for the harmonic oscillation model in section 5. We approximate the movement of the COM during walking or running by a Duffing equation, i.e. a homogeneous non-linear second order differential equation. We write

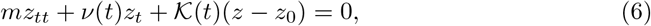

where, unlike the harmonic oscillation model from Section 5, we consider stiffness and viscosity to not be constant, but functions depending on time, 𝒦 (*t*) and *v*(*t*). This makes sense, as stiffness and viscosity depend on a phase of a stride. For example, the slope on Figures 2 and 3 is changing, and the slope in the plane *z, z*_*tt*_ reflects stiffness: it is 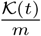 at a time *t*. We divide both sides by *m* and consider the approximation of the functions 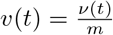 and 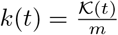 by cubic polynomials,

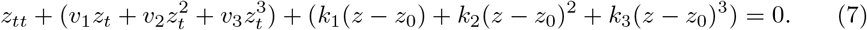

As the value *z*_0_ is not known beforehand, we look for the best fitting curves of the form of least squares:

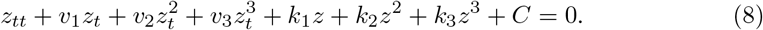

The best fitting curve in the chosen coordinates is a differential equation of second order. We visualize the solution of this differential equation as a curve in the same coordinate system as the data curve. See, for example, Figures 4, 7, 8, where the red curve is the data curve, and the blue curve is the solution of the differential equation given by the best fitting curve. Initial conditions are the values *z*(0) and *z*_*t*_(0). Equation 8 is equivalent to Eq. (7) after expanding brackets and renaming coefficients. From our data, the viscosity coefficients *v*_1_, *v*_2_, *v*_3_ turned out to be zero if the FFT threshold is large enough, for instance if it is *≥*0.3. This is not surprising because we have assumed no excitation force (homogeneous equation), and the participants have maintained stable walking/running. If the viscosity is non-zero, the curve is a stable spiral approaching *z*_0_, as in Fig. 8. Scaled mean squared error between the Duffing equation output and the observed data was computed. As the values *z, z*_*t*_ and *z*_*tt*_ have different units, each value was scaled by dividing by the difference between the minimum and maximum values on the corresponding axis.

**Fig 4.**
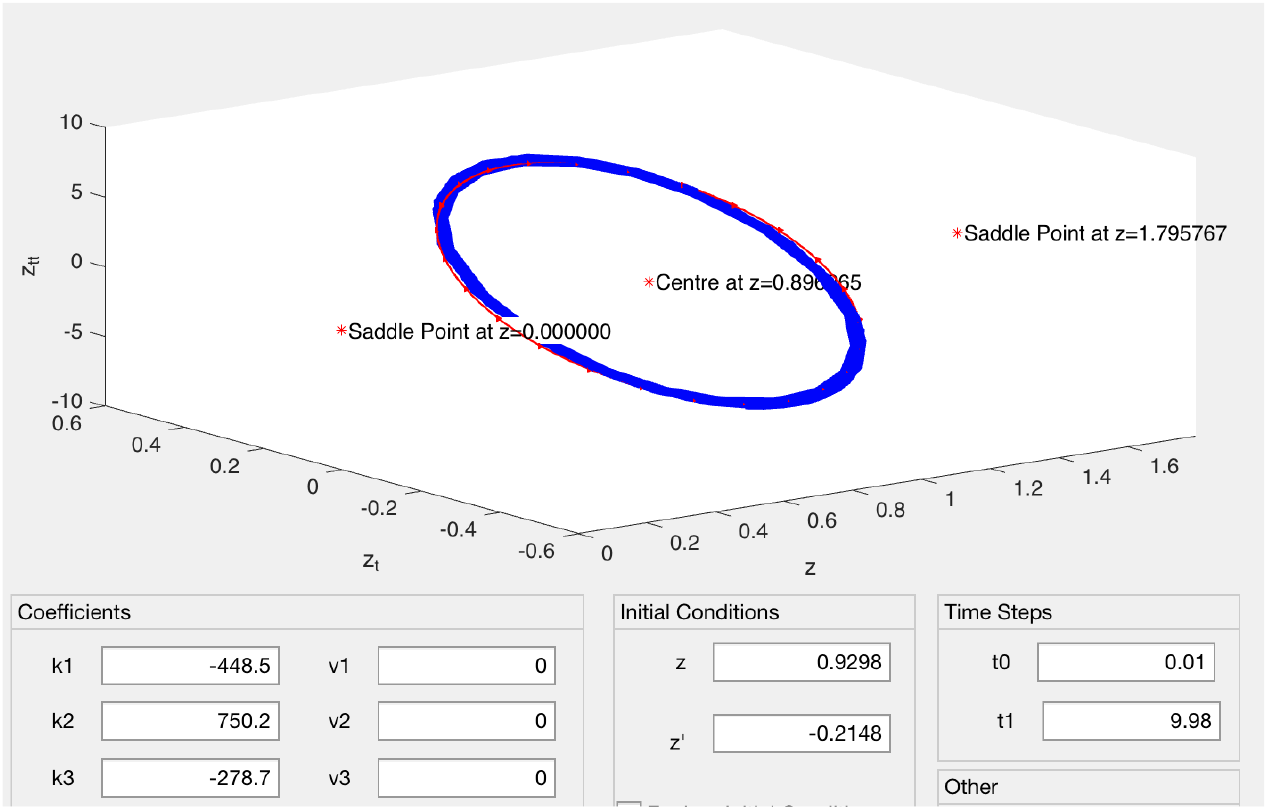
Curve (blue) for the solution to Eq. (8) with coefficients *k*_1_, *k*_2_, and *k*_3_ = *−k* for a participant (178cm tall, running at 8 km/h) compared with the data curve (red). The mean squared error is 0.015. The fixed points are saddles at *z* = 0 and *z ≈* 1.8 and the centre at *z*_0_ *≈* 0.9

**Fig 5.**
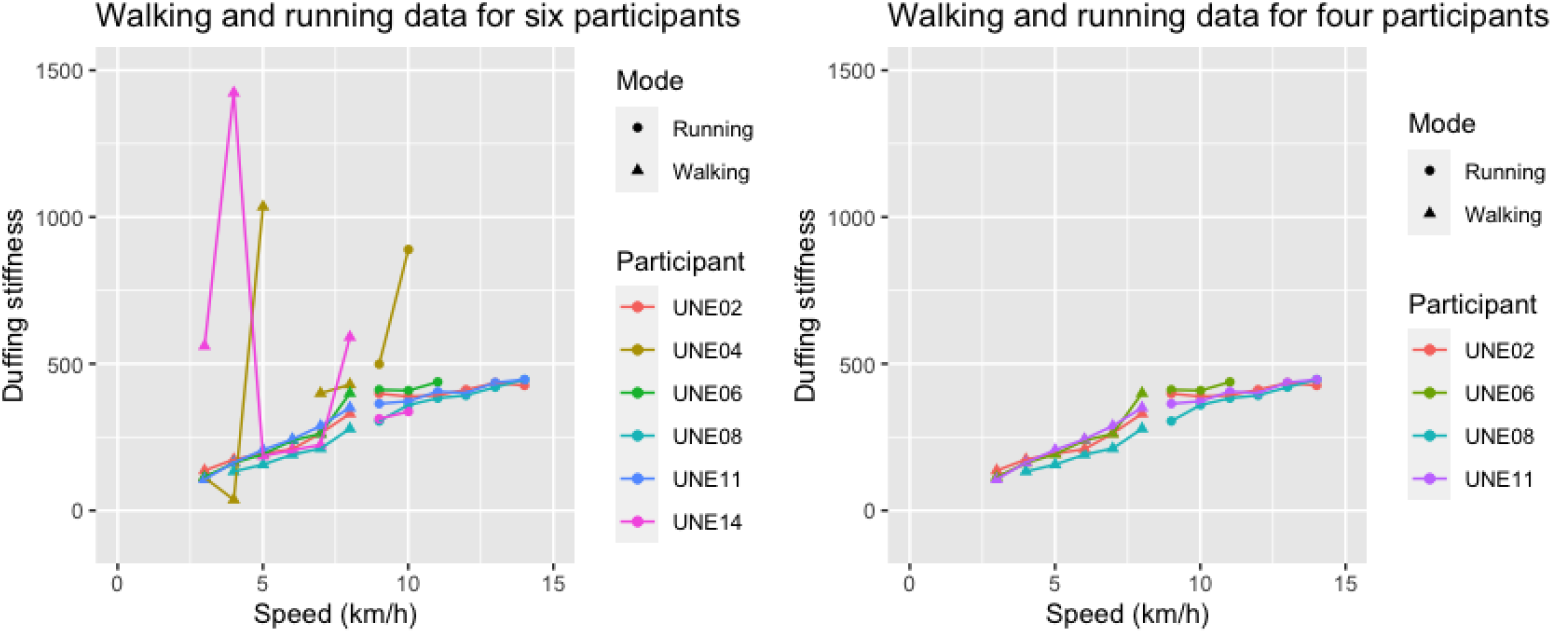
Duffing stiffness depending on speed with and without outliers. We consider the models for walking and for running separately

### 6.2 Finding and analysing the fixed points

We are also interested in the fixed (equilibrium) points [16] for the curves described by Eq. (8). We first rewrite the second order differential Eq. (7) as a system of first order differential equations:

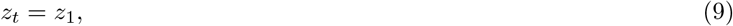

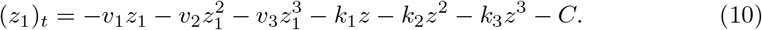

We remind that for stable gait *v*_1_ = *v*_2_ = *v*_3_ = 0. The fixed points are found by solving the equilibrium equations (see for example [3])

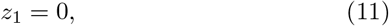

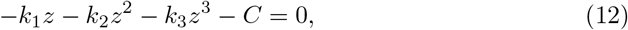

i.e. the fixed points are the roots of the cubic equation *k*_1_*z* + *k*_2_*z*^2^ + *k*_3_*z*^3^ + *C* = 0. The solutions to Eq. (8) are plotted in the same coordinate system as the data curves, see for example Figure 4. The solution curves are stable if we set one fixed point equal to zero, i.e. *C* = 0. Then the two other fixed points occur at *z*_0_ (the centre of the closed curve, average coordinate of the centre of mass during walking/running), and at *h*, which is approximately 2*z*_0_ for stable walking/running. The differential equation Eq. (8) becomes

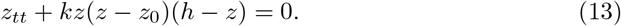

The connection between the coefficients of Eq. (8) and Eq. 13 is

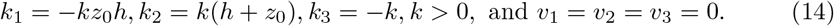

Numerical computations show that, for stable gait, the fixed point *z* = *z*_0_ is a centre, while the fixed points *z* = 0 and *z* = *h* are saddles.

### 6.3 Interpretation of the parameters in the model

Eq. (13) shows that we can model the movement of the centre of mass by a Duffing equation, up to a constant *k*, knowing only *h* and *z*_0_. The values *h* and *z*_0_ are close to the height of a person and to the average coordinate of the COM in motion, correspondingly. The gravitation constant *g* is involved implicitly in the differential equation, in a similar way as in the harmonic oscillation model in Section 5.

The coefficient *k* does not have the meaning of the square of the angular frequency, 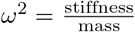, as in a linear case (Section 5), but *k* behaves in a similar way: it increases with an increase of the walking/running speed. We call this constant the **Duffing stiffness**. The meaning of the coefficient *k* is found from the following consideration. We rewrite Eq. (13) as

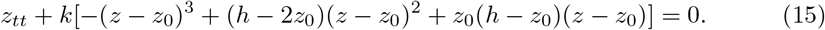

For *z* close to *z*_0_ the linear approximation at *z*_0_ is

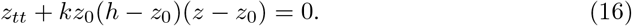

Comparing this equation with Hook’s Law (Eq. (4)) we get an expression that relates the angular frequency *ω* with the coefficient *k* and the values *z*_0_ and *h*: *ω*^2^ *≈ kz*_0_(*h − z*_0_). Hence, the Duffing stiffness

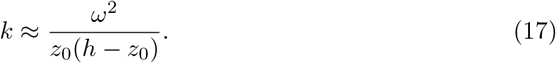

### 6.4 Case *h* = 2*z*_0_

Assuming that the centre *z*_0_ is approximately in the middle between two saddle points, 0 and *h*, we get *h* = 2*z*_0_, and Eq. (15) becomes

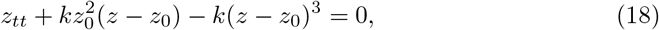

where *k >* 0, and

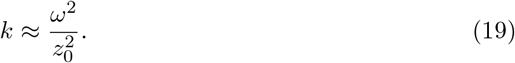

Eq. (18) is the Duffing equation for a softening oscillator [3], i.e. the stiffness diminishes with the displacement.

### 6.5 Example based on the collected data

Figure 4 shows the observed smoothed data curve (red) and the curve corresponding to the solution of Eq. (13) (blue) for one of the participants, together with the fixed points. The differential equation that describes the movement of the centre of mass of this participant running at the speed 8 km/hr is *z*_*tt*_ + 279(*z −* 0.9)(1.8 *− z*) = 0. We have rounded |*k*_3_| = *k* to an integer, and *z*_0_ and *h* to the first decimal place. See Figure 4.

We can check that Eq. (4) and the approximation Eq. (17) are realistic by computing the number of strides in a second and comparing it with the real value.

From *k* = 279, 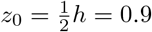 we can compute 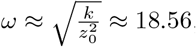, i.e. the number of strides in a second is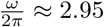. This participant made 24 strides in 10 secs, i.e. the real number of strides in a second is 2.4.

## 7 Statistical Analysis

### 7.1 UNE data

Fig.5 (left) shows that the Duffing stiffness depends on the speed of walking/running for each participant. By visually inspecting the graph, we see some outlier points. To further examine these, we calculated the standard deviation of Duffing stiffness for each participant, for the walking data. Two participants had standard deviations *>* 400, while the rest of the participants had standard deviations *<* 100. Our observation of these two participants during the data recording showed that they were uncomfortable with some of the speeds, and they reported having no experience with treadmills. Therefore, we excluded these two participants from all further analyses. Fig.5 (right) shows the data for the remaining four participants.

The behaviour of the Duffing stiffness is different for walking (3 *−* 8 *km/hr*) and running (9 *−* 14 *km/hr*). Therefore, we separately fit data for walking and running speeds. We fitted a Linear Mixed Effect model for the walking data. We included speed as the fixed effect and Duffing stiffness as the dependent variable. We allowed both the slopes and the intercept to vary across participants. The model showed a slope estimate of 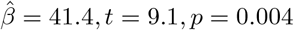 . The intercept was −14.6, *p* = 0.5.

For the running data, we performed an equivalent analysis. Here, the slope associated with speed was not significant, though trending in a positive direction, 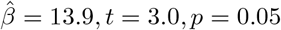. The intercept was 245.6, *t* = 4.0, *p* = 0.02.

### 7.2 Comparing UNE dataset against public datasets

Next, we compare our data against public datasets for 42 walking participants [10], and 29 running participants [11]. As these sources were set up slightly differently than the UNE data, in order to match the data, we estimated the COM during walking or running, using the midpoint of the ASIS markers and subtracting the lowest value for the heels marker.

Figure 6 shows the relationship between speed and Duffing stiffness.

**Fig 6.**
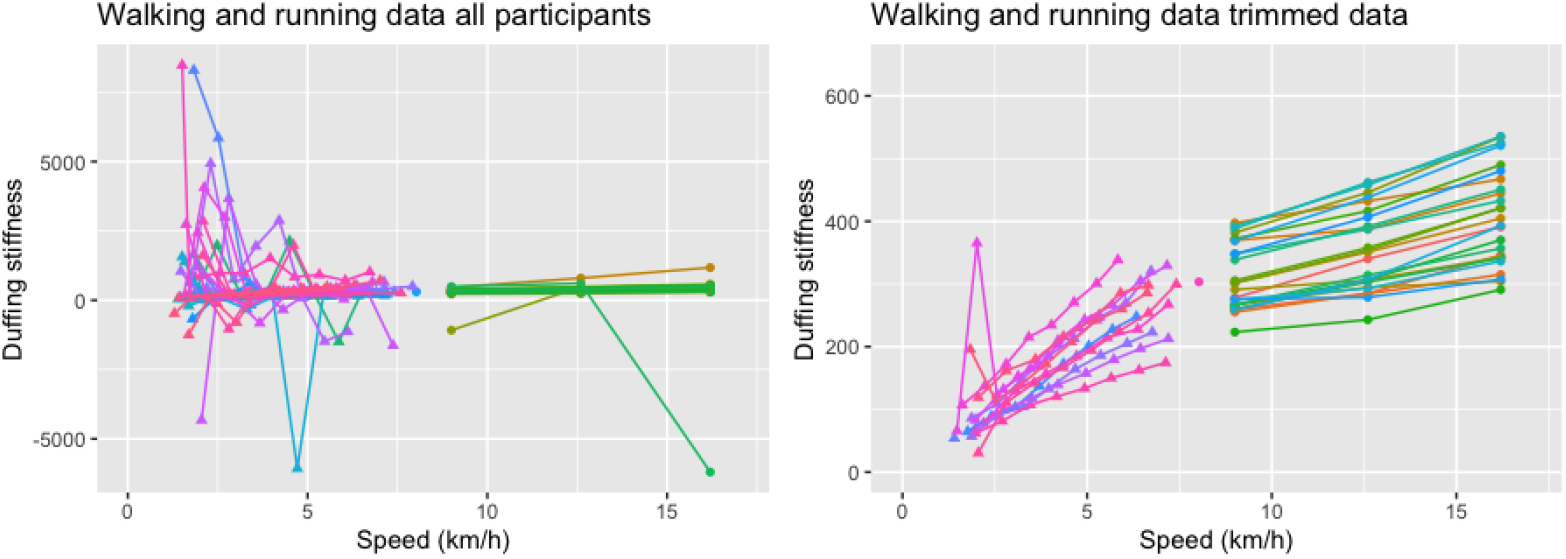
Duffing stiffness as a function of speed, with and without outliers. Note that the Duffing stiffness, *k*, depends also on the height of the COM of a participant, *z*_0_, and the angular frequency of walking/running, 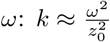

**Fig 7.**
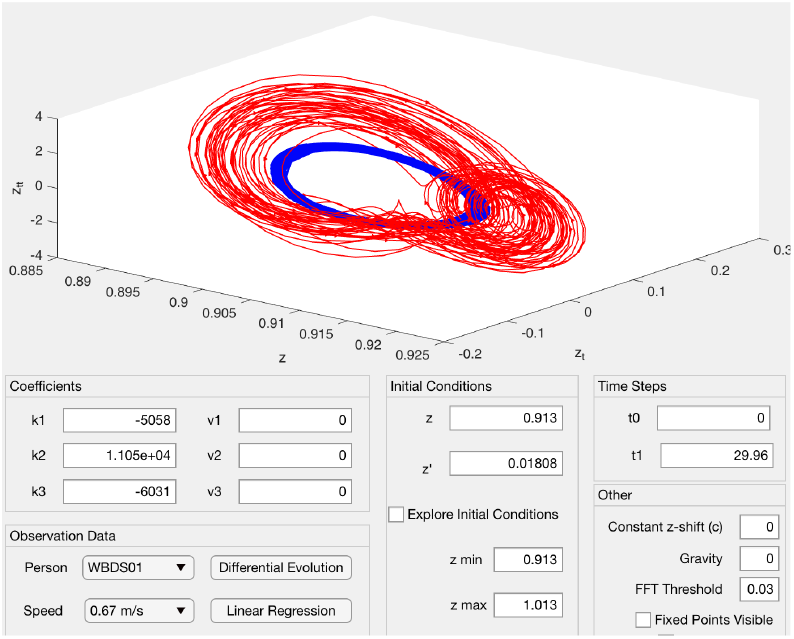
Slow walking speeds (*<* 3 *km/hr*)

**Fig 8.**
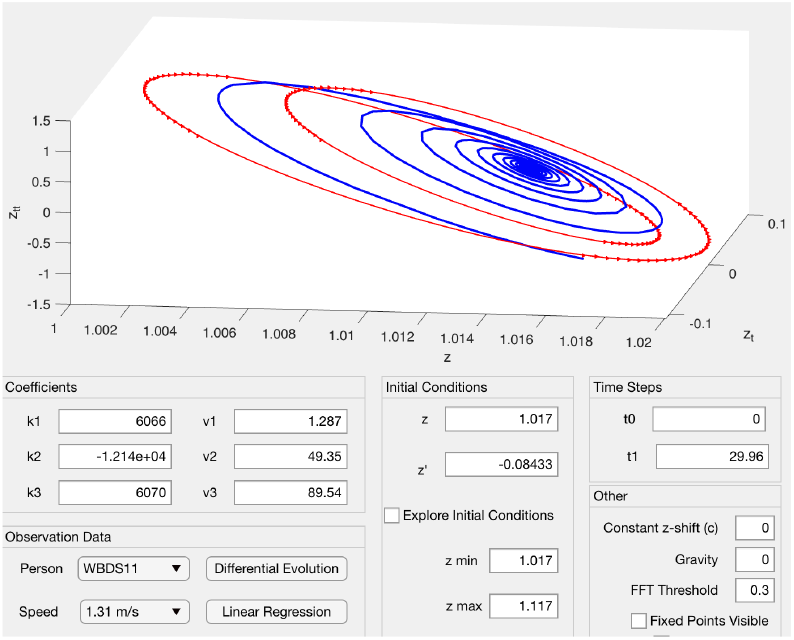
Asymmetry of the gait due to often contain an additional loop because of different strengths of the left and the right the compelled braking during each step legs

First we removed one participant with only one data point. Then, to identify outliers, we calculated the standard deviation in Duffing stiffness for each participant and excluded outliers for the walking and the running datasets, separately. For the walking data, [11], the standard deviation ranged from 39.2 to 3183.5. We chose to remove all participants with a standard deviation *>* 100. This left us with 12 participants for the walking data. We proceeded to fit the data with a Linear Mixed Effect model, akin to the UNE data. Again, we get a significant effect of speed on Duffing stiffness, 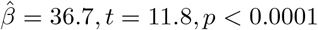. The intercept is not significant, 24.0, *p* = 0.09. For the running data, [10], we removed three out of 29 participants with standard deviations *>* 100. The Linear Mixed Effect model showed a significant effect of speed, 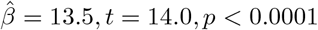. The intercept is 188.1, *t* = 19.7, *p <* 0.0001.

Finally, we compared the two datasets against each other, to determine whether there are any overall differences in Duffing stiffness, or whether there is difference in the relationship between speed and Duffing stiffness. We created two models, one for running and one for walking, including both of the trimmed datasets. In the Linear Mixed Effect model, Duffing stiffness acted as the dependent variable, and the fixed effects were speed, dataset UNE versus [10, 11], with UNE acting as a baseline. We allowed the slope and the intercept to vary across participants. For walking, the effect of speed was, again, significant, 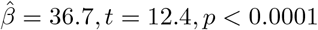. However, the effect of dataset was not significant, 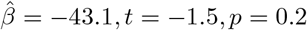, nor was the interaction between dataset and speed, 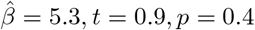. Similarly, in the running dataset, the effect of speed was 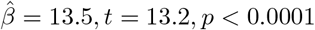. Neither the effect of dataset, nor the interaction of dataset and speed were significant, 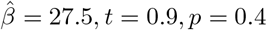 and 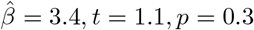, respectively.

In summary, we found, overall, a significant positive relationship between speed and Duffing stiffness. For the running UNE data, the slope was not significant; however, when we combined the two datasets, the running slope was significant, and we found no interaction. Thus, the lack of significance in the UNE running data may be a result of low statistical power. We found no main effect of dataset, nor an interaction between dataset and speed. Thus, we find no evidence of a difference across datasets.

### 7.3 Identifying and examining outliers

From a practical perspective, an interesting aspect is participants whose data deviates from the fitted model. Here, we defined outliers based on standard deviation. This is the simplest method, which can easily be applied by a sport scientist. In our analyses, we drew a somewhat arbitrary threshold, where we treated all participants with a standard deviation *>* 100 as outliers. In practice, this can act as a sign for the practitioner to follow up on the walker or runner if their standard deviation is high. The reasons for high standard deviations could be different. For example, a typical problem for uncomfortable speeds less than 3 km/hr is an additional loop, as on the red line in Fig.7. The Duffing equation does not take into account the loop, as the blue approximation curve demonstrates. The other example of an outlier is illustrated in Fig. 8, where the curve brakes down into two parts, corresponding to the left or to the right leg. Red lines represent the data, blue lines are the solutions of the differential equation

A third reason for outliers that we noticed is an instability of walk, when each step varies in the amplitude and in the average height.

## 8 Duffing equation with viscosity and excitation force

### 8.1 Motivation

Approximation by a homogeneous equation with zero viscosity does not take into account the asymmetry caused by damping and excitation forces.

For example, Figure 9 shows a smoothed data curve for a running participant (14 km/hr, FFT threshold=0.3) in the phase plane (*z, z*_*t*_). This curve is close to an ellipse, and the absolute value of the slope of the main axis of the ellipse, CA, is equal to 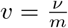 if we assume that the viscosity is a constant. (If the viscosity *ν* = 0, the slope is vanishing, and the corresponding axis is horizontal.) The symmetry with respect to *z*_*t*_ = 0 is disturbed by the slope.

**Fig 9.**
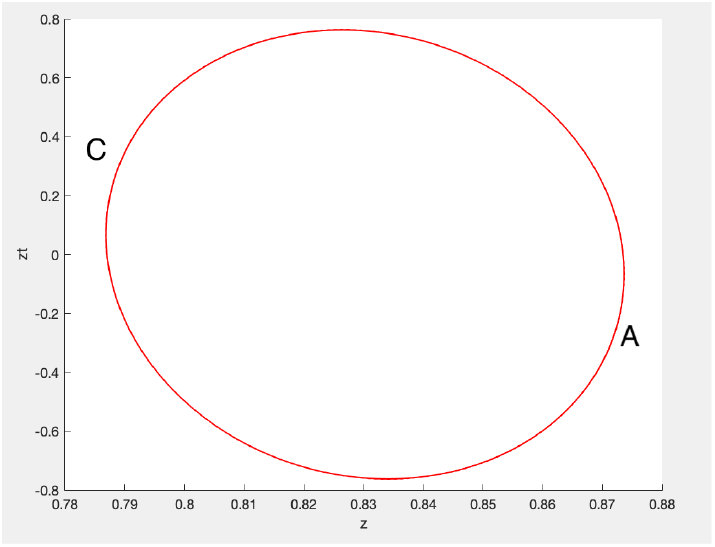
The absolute value of the slope MN multiplied by mass *m* is the viscosity *v*

Now we compare this slightly asymmetrical typical data curve with the symmetrical energy level curves, see Figure 10. Solutions of Eq. (13) with different initial values of *z* (*z*_*t*_ = 0) give a set of energy level curves. The red curve is the data curve, the movement occurs in the direction ABCD. Energy is gained twice in each cycle (gait) in a phase of a “step”, BC, and in a phase of a “fall” (in general not free fall), DA. The maximum of the energy occurs at points A and C, the minimum occurs when at points B and D. There is a natural desire to find an approximation that considers the viscosity and the restoring force.

**Fig 10.**
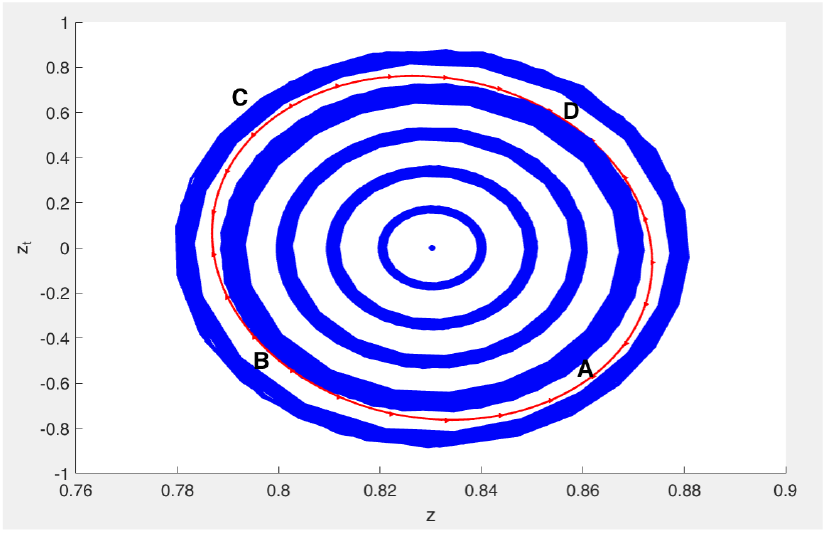
The energy level curves (blue) and the data curve (red), the same as in Fig. 9, in the phase plane (*z, z*_*t*_)

### 8.2 Looking for the best fitting curve for a non homogeneous differential equation

We were looking for the best fitting curve in different forms, as for example *z*_*tt*_ + *vz*_*t*_ + *C*_1_*z* + *C*_2_*z*^2^ + *C*_3_*z*^3^ = *f* cos(Ω*t − ϕ*), where the term *vz*_*t*_ is the linear approximation of the damping force divided by mass, *C*_1_, *C*_2_, and *C*_3_ are coefficients of the cubic nonlinearity. The constant 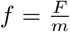, where *F* is the amplitude of the excitation force and *m* is the mass, Ω is the angular frequency of the excitation force. The gravitation constant *g* is involved in the equation in a similar way as for the two already discussed models. We have also tried to find the best fitting curve in the forms based on Eqs. (13),(18). As in the previous models, the equation for the best fitting curve was interpreted as a second order differential equation. The solution of the equation was computed and plotted in the same system of coordinates as the data curve. However, the balance between the viscosity and excitation force was too delicate, and most solution curves became stable spirals. The balance worked with an FFT threshold of 0.5, at least for some curves, based on Eq. (18),

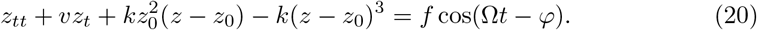

### 8.3 Example

In Figure 11 the red data curve is approximated by a homogeneous differential Eq. (13) (blue curve), and by Eq. (20) containing the damping and restoring forces (green curve). The mean squared error for the homogeneous equation is 0.01. The non-homogeneous equation gives a better approximation with the mean squared error 0.002.

**Fig 11.**
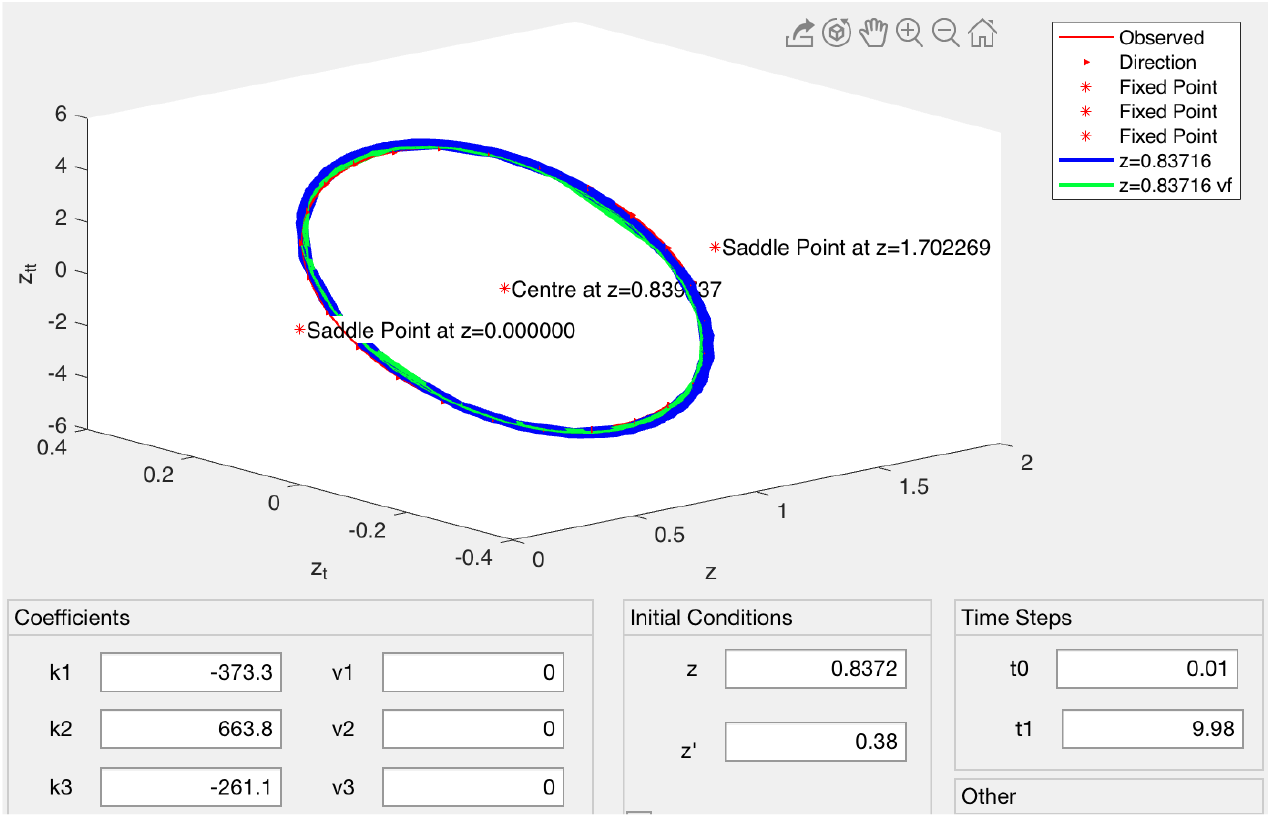
The data curve (red) is approximated by a second order homogeneous differential equation (blue) with the mean squared error 0.01, and by a second order non-homogeneous differential equation (green) with the mean squared error 0.002

## 9 Plans for future research

### 9.1 Classification of curves

One of the ideas for future research is to compare and classify curves for different participants. The curves can have individual features, see for example Figure 12 (FFT threshold 0.03).

**Fig 12.**
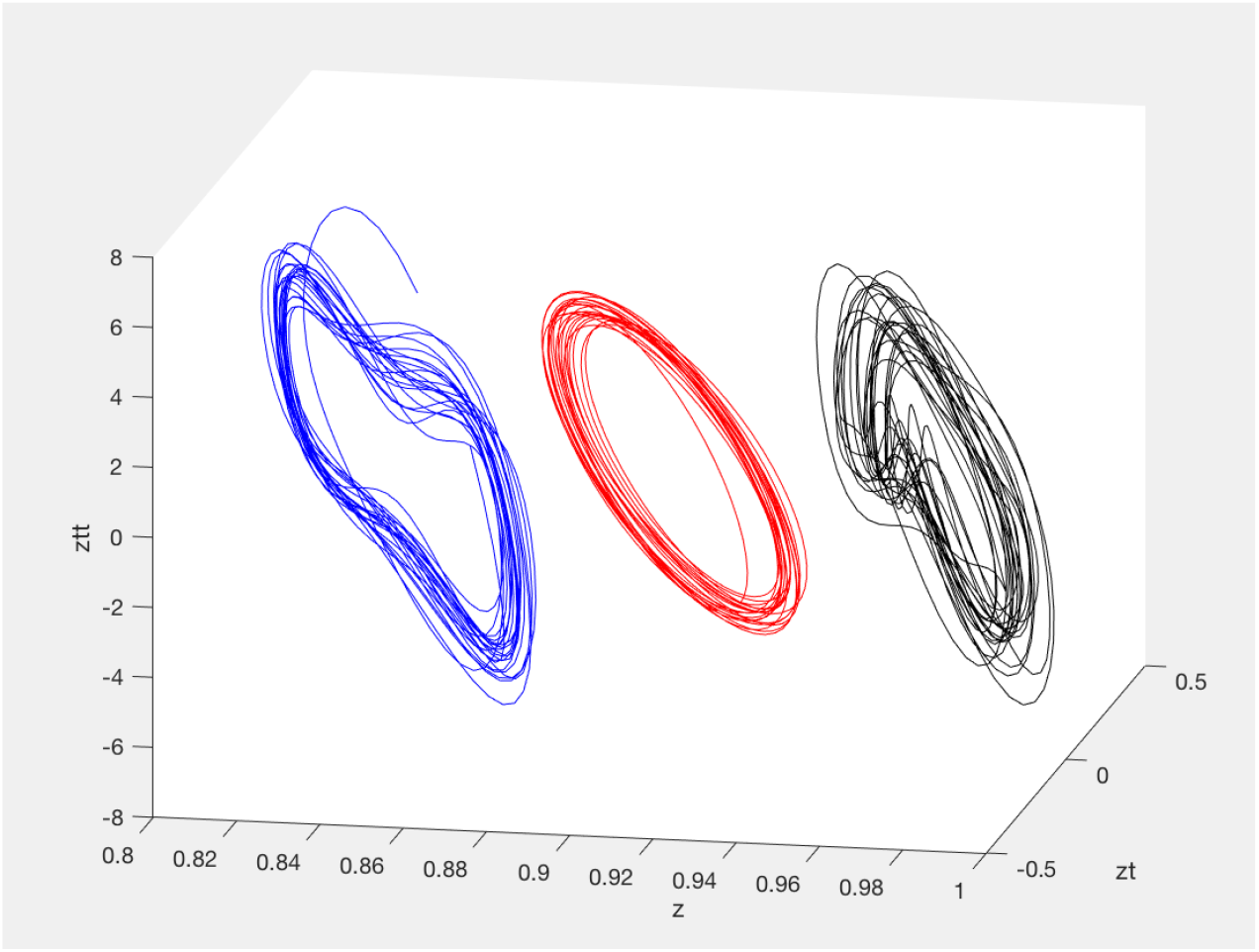
Comparison of curves for three participants walking at 6 km/hr

Also the left and right leg strides are almost indistinguishable if we chose an FFT threshold *≥*0.3. But for some participants, we observe a difference in the slopes for even and odd strides if the FFT threshold is smaller, suggesting sensitivity in identification of gait asymmetries. In Figure 13 the green curve (FFT threshold = 0.03) is split into two parts. Recall that different slopes correspond to different angular frequencies. The black curve shows the same data smoothed with an FFT threshold of 0.3. Bipedal models were investigated using a rather mechanical approach in several papers as for example in [12, 22].

**Fig 13.**
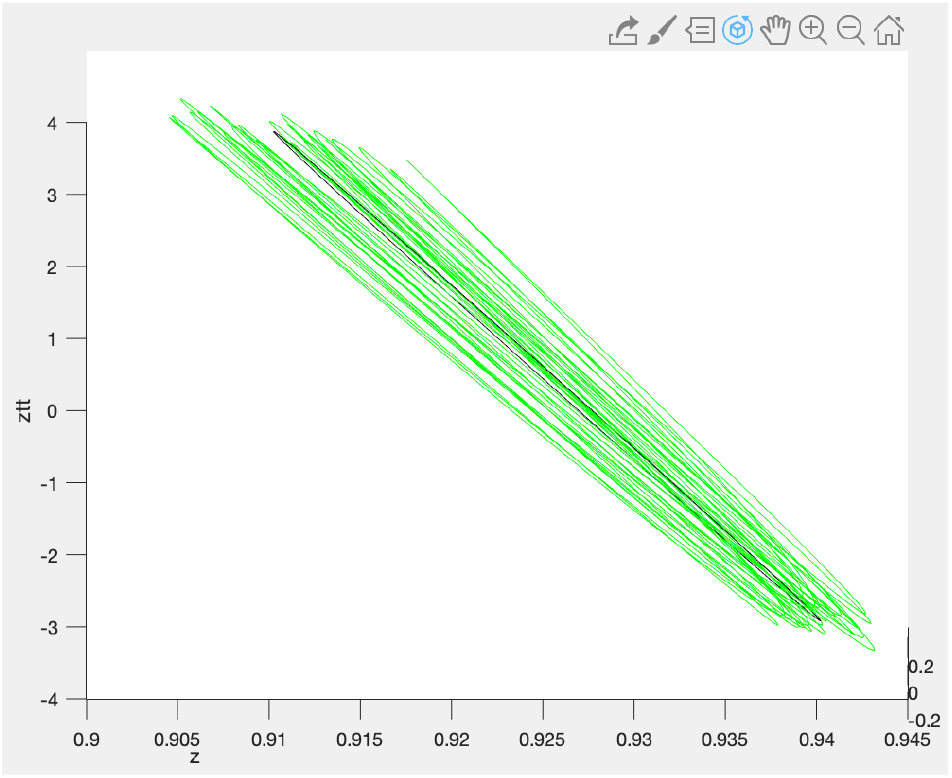
Two curves for a participant walking 6 km/hr. The green curve for FFT threshold 0.03, the black curve for FFT threshold 0.3

### 9.2 Horizontal components

In this paper we have modelled only the vertical component of the movement of the COM. We plan to also investigate the horizontal components in the future. Our hypothesis is that the horizontal component that is perpendicular to the forward movement can be approximated by a second order differential equation with negative stiffness.

### 9.3 Investigating possible reasons for injuries

Our model gives an insight into the dynamics of human gait. The Duffing stiffness at a particular speed is approximately the same for each person who maintains a stable gait. If the Duffing stiffness differs, it points to some uncomfortableness in walking or running. The reason for this uncomfortableness can be investigated by looking at the curves closely. We plan to investigate possible reasons for injuries by determining how the stress is generated. One possible idea is to investigate why female runners have more frequent ACL (anterior cruciate ligament) tears than men [27].

## 10 Conclusion

We have introduced an innovative method for the investigation of human gait. This method is based on the visualisation of the vertical component of the movement of the centre of mass during walking or running, in the space of the coordinates position, velocity, and acceleration of the centre of mass. This method allows us to compute leg stiffness for an established harmonic oscillator model in a new way, as the slope of the best fitting plane. We determined some differences and some common features for walking and running curves. In this paper we concentrated on features that are common for walking and for running. We suggested a model by a non-linear homogeneous differential equation. For this model we need to know the average of the COM, *z*_0_, during the gait motion, and a coefficient *K* that depends on a participant and on a walking/running speed. Assuming *h* = 2*z*_0_, we get the Duffing equation for a softening oscillator

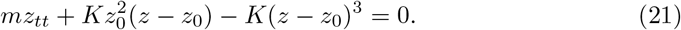

Statistics showing the dependence of the Duffing stiffness on the speed was shown separately for walking and for running speeds, using both our collected dataset and an open dataset [10, 11].

We also had a partial success in approximation of the movement by a second order non-linear non-homogeneous equation

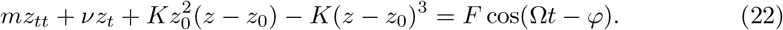

We believe that our new method is a tool that could lead to other interesting and novel results.

## 11 Acknowledgements

The study followed ethical protocols as per ethics requirements (HE19-239). Thanks to computer science students Ben Fisk, Danielle Galvin and Jarra McIntyre for developing a prototype of the software used for this paper and to sport science student Megan Bancks for the literature research. Thanks to Adam Harris and Gerd Schmalz for their critical comments and support.

## Notes

https://doi.org/10.6084/m9.figshare.15041322.v2

https://github.com/davidjohnpaul/une_gait

### Competing Interest Statement

The authors have declared no competing interest.

